# Microscopy-Guided Spatial Proteomics Reveals Novel Proteins at the Mitochondria-Lipid Droplet Interface and Their Role in Lipid Metabolism

**DOI:** 10.1101/2025.01.24.626799

**Authors:** Yen-Ming Lin, Weng Man Chong, Chun-Kai Huang, Hsiao-Jen Chang, Chantal Hoi Yin Cheung, Jung-Chi Liao

## Abstract

Mitochondria-lipid droplet (LD) interactions play a critical role in lipid metabolism and the progression of metabolic diseases such as non-alcoholic fatty liver disease (NAFLD). However, the dynamic nature of these interactions has hindered the identification of novel protein constituents and their functional roles. Here, we employed Microscoop Mint, an advanced microscopy-guided spatial proteomics platform, to isolate proteins localized at mitochondria-LD contact sites in oleic acid (OA)-treated HepG2 cells, an in vitro model for studying fatty liver disease. Microscoop Mint integrates high-resolution image analysis and two-photon illumination to achieve precise biotinylation of proteins at subcellular regions of interest. Coupled with mass spectrometry, this approach identified a proteome enriched at the mitochondria-LD interface, including both well-characterized lipid-associated proteins and previously unrecognized candidates. Among the 373 common proteins identified across replicates, five novel candidates with no prior association to lipid metabolism were selected for validation. Immunofluorescence staining confirmed their localization at mitochondria-LD contact sites, with more pronounced association in OA-treated cells. Notably, suppression of one candidate, FHL3, led to reduced LD size, elongated mitochondrial morphology, and diminished mitochondria-LD interactions, suggesting its role in regulating fatty acid β-oxidation. Our findings demonstrate the utility of Microscoop Mint in unveiling novel molecular players at organelle contact sites. This study not only provides insights into the functional dynamics of mitochondria-LD interactions but also highlights potential therapeutic targets for lipid metabolic disorders and NAFLD.

## Introduction

Membrane-bound organelles are a hallmark of eukaryotic cells, providing spatio-temporal compartmentalization for diverse biochemical processes. Communication and coordination among organelles are crucial for maintaining cellular homeostasis, facilitated by membrane contact sites (MCSs) that serve as hubs for metabolite exchange (1). Among these organelles, lipid droplets (LDs), which are composed of a phospholipid monolayer surrounding a core of neutral lipid insides, act as energy storage depots and engage in dynamic interactions with other organelles (2), particularly mitochondria (3). Mitochondria are central to lipid transfer and metabolism, including fatty acid β-oxidation for energy production (4). Disruption of mitochondria-LD communication or mitochondrial dysfunction can lead to metabolic disorders, including non-alcoholic fatty liver disease (NAFLD), also referred to as metabolic-associated fatty liver disease (MAFLD) (5). NAFLD is a chronic, progressive condition characterized by hepatic steatosis, which is unrelated to excessive alcohol intake and other liver damage. The “multiple hit” hypothesis has emerged to explain the complex pathogenesis of NAFLD (6) highlighting factors such as increased lipolysis in visceral adipose tissue (AT), enhanced hepatic de novo lipogenesis (DNL), and high-calorie or high-fat diets, all of which contribute to hepatic lipid accumulation in the liver (6). Furthermore, impaired mitochondrial β-oxidation, the primary pathway for fatty acid degradation, along with elevated DNL, further exacerbate lipid accumulation, driving the progression of NAFLD (7, 8).

Saturated palmitic acid (PA) and monounsaturated oleic acid (OA) are the predominant fatty acid found in steatotic liver (9). These fatty acids are mainly derived from dietary sources, absorbed in the intestine, esterified into triglycerides (TG), and transported to the liver. Excessive accumulation of PA or OA leads to TG storage within lipid droplets in liver cells (10). While PA overload is associated with mitochondrial dysfunction and liver damage, OA has been shown to protect cells from lipotoxicity (11). Notably, studies indicate that OA overload enhances LD-mitochondria interactions and promotes LD growth (12). However, the mechanisms underlying OA’s protective effects remain unclear.

Mitochondria are central to energy metabolism, and their dysfunction has been linked to the development of fatty liver disease, highlighting their critical role in the pathogenesis of NAFLD. Recent studies emphasize the importance of peri-droplet mitochondria (PDM) in maintaining lipid homeostasis. Key proteins such as perilipin-5 (PLIN5), intermembrane lipid transfer protein VPS13D (VPS13D) and synaptosomal-associated protein 23 (SANP23) have been identified as essential for maintaining the structural and functional integrity of PDM (13). These LD-associated proteins were either transferred from the endoplasmic reticulum (ER) or recruited from the cytoplasm through amphipathic helices (14, 15), where they play pivotal roles in regulating lipid metabolism. Despite advances in proteomic analyses of isolated LDs, the dynamic and complex nature of mitochondria-LD interactions continues to pose significant challenges in identifying novel protein components and elucidating their roles in mitochondria-LD communication.

Spatial proteomics technologies offer greater scalability and deliver valuable insights into cellular functions (16). To address these challenges, we employed our developed Microscoop Mint (17), a novel microscopy-guided spatial proteomic platform that enables precise biotin tagging of subcellular regions of interest with a resolution of 240 nm. By integrating Microscoop with LC-MS/MS, we identified the proteome at these specified regions with high precision. Using this innovative approach and in vitro steatosis model, we conducted a comprehensive analysis of mitochondria-LD-associated proteins in OA-overloaded HepG2 cells. This study uncovered proteins involved in regulating mitochondria-LD interactions, shedding light on the potential protective role of OA against lipotoxicity.

Mass spectrometry analysis following Microscoop labeling identified the proteome of the mitochondria-LD interface, successfully recovering known proteins previously implicated in mitochondria-LD interactions, such as perilipin-2 (PLIN2), alpha/beta hydrolase domain-containing protein 5 (ABHD5), and patatin-like phospholipase domain-containing protein 2 (PNPLA2/ATGL). Beyond these well-characterized LD-associated proteins, we identified 14 novel high rank proteins with no prior association to LDs. Among these, five candidates, including a mitochondrion protein with the highest fold change, were selected for further characterization. Intriguingly, immunofluorescence staining confirmed their presence at mitochondria-LD contact sites and around lipid droplets, particularly in OA-treated HepG2 cells compared to vehicle-treated cells. These novel proteins are known to participate in diverse biological processes such as growth regulation, proliferation, transcription, and splicing, yet have not been linked to lipid metabolism, suggesting potentially new roles in mitochondria-LD communication under conditions of LD accumulation.

Notably, suppression of one candidate, four-and-a-half LIM-only protein 3 (FHL3), reduced mitochondria-LD contact, decreased LD size, and led to elongated mitochondria in OA-treated FHL3-deficiency HepG2 cells. Mitochondrial elongation suggests a reduction in fatty acid β-oxidation (18), indicating that FHL3 may play a critical role in regulating mitochondria-LD communication and lipid metabolism processes. In summary, through Microscoop’s spatial protein purification capabilities, our study identifies previously unrecognized protein constituents at the mitochondria-LD interface, offering new avenues for investigating lipid regulation and its connection to NAFLD pathogenesis.

## Materials and methods

### Cell culture, reagents, and transfection

HepG2 cells were cultured in DMEM/F12 medium (11320-033, Gibco, Thermo Fisher Scientific, Waltham, MA) containing 10% FBS (A52567-01, Gibco, Thermo Fisher Scientific, Waltham, MA), antibiotics (15140-122, Gibco, Thermo Fisher Scientific, Waltham, MA), and maintained in a 37℃ humidified incubator with 5% CO_2_. Lipid droplet accumulation was induced by treating cells with 200 μM BSA-complexed oleic acid (molar ratio of FFA: BSA is 5:1; SI-O1383, SI-A8806, Sigma-Aldrich, St. Louis, MO) for 16 h. Lipid droplet were subsequently stained using 1 μM BODIPY 558/568 (D3835, Thermo Fisher Scientific, Waltham, MA) for 1 h. For gene silencing experiments, the scramble and two FHL3 siRNAs (s223535, AUCUAGACUUCUCUUUAUA, and s5200, CGAGAAUGUCUGGUCUGUA) were purchased from Invitrogen (Carlsbad, CA). Transfection was performed using the reverse transfection method with GenMute siRNA transfection reagent (SL100568-HEPG2, SignaGen Laboratories, Frederick, MD), following the manufacturer’s protocol. Equal amounts of scramble or FHL3 siRNA were diluted in transfection buffer (SL100572, SignaGen Laboratories, Frederick, MD), and a corresponding volume of transfection reagent was added to the diluted siRNA. The mixture was gently mixed and incubated at room temperature for 15 min. The transfection complex was then added to HepG2 cells in suspension, mixed by gently rocking and incubated for 72 h before further analysis.

### Antibodies

The antibodies used in this study are listed in Supplementary Table 1.

### Immunofluorescence staining

HepG2 Cells were seeded at approximately 70% confluency in chambered cover glass (C1-1.5H-N, Cellvis, Mountain View, CA) one day prior, then fixed with 2.4% paraformaldehyde for 5 min and permeabilized with cold methanol for an additional 3 min. Following three washes with PBS, the cells were blocked for 1 h at room temperature using 3% BSA diluted in PBS-Tx (PBS containing 0.1% Triton X-100) and then washed three times with PBS. For Microscoop photolabeling, blocking was performed sequentially using Block 1 and Block 2 blocking reagents from the Synlight-Rich^TM^ kit (SYN-RI0106, Syncell, Taiwan) for 1 h and 30 min, respectively. Cells were incubated with corresponding primary antibodies overnight at 4℃, followed by a 1 h incubation with fluorescence probe-conjugated secondary antibodies. To validate biotinylated proteins validation after photolabeling, cells were stained with verifying reagent from the Synlight-Rich^TM^ kit (SYN-RI0106, Syncell, Taiwan) for 1 h at room temperature. After three PBS washes, cell nuclei were stained with Hoechst-33342 (H3570, Thermo Fisher Scientific, Waltham, MA). Images were captured using either epi-fluorescence microscope (TE2000, NIKON, Japan) or confocal microscope (LSM880, ZEISS, Germany).

### Microscoop photolabeling

For each replicate, 8 chambered cover glass containing IF stained HepG2 cells were prepared and evenly divided into two groups - one designated for photolabeling with Microscoop Mint (MS.1012, Syncell, Taiwan), while the other served as an unlabeled control. The photoreactive probe (Photolabel) from Synlight-Rich^TM^ kit was added to the chambered cover glass of both groups according following the manufacturer’s instruction. Images acquisition and mask generation for mitochondria (detected by Tom20 antibody) and lipid droplet membrane (detected using PLIN2 antibody) were conducted using the Autoscoop^TM^ software. First, the ‘Contrast Limit Equalization’ function was applied to adjust contrast, brightness and enhance edge definitions in both images. The ‘Adaptive Threshold’ function was then used to convert gray scale image into binary images, allowing the definition of regions of interest (ROIs) for mitochondria and lipid droplet membranes based on intensity. Masks for each were generated accordingly. The ‘Size Filter’ function was applied to filter out small noise particles from the masks. To identify mitochondria-LD contact sites, the ‘Mask And’ function was applied to overlap the mask of mitochondria and lipid droplet membranes, defining the intersecting regions as contact sites. For each field of view, these image processing steps were applied automatically, marking mitochondria-LD contact sites. A laser was then precisely illuminated on these specific regions to initiate biotinylation. The photolabeling process was repeated across the entire slide automatically, ensuring complete labeling of the slide. After completing photolabeling, the photoreactive probes were removed, and the cells were thoroughly washed with PBS-Tx to minimize interference from unreacted probes during subsequent pulldown procedures.

### LC-MS/MS analysis

The samples for mass spectrometry were prepared using the Synpull^TM^ reagent kit (SYN-PU0106, Syncell, Taiwan) according to the manufacturer’s instruction. Briefly, photolabeled and unlabeled cells were scraped and collected into separate tubes using scraping buffer (solution A). The samples were centrifuged at 5000 x g for 3 min at 4℃, and the supernatant was removed, leaving approximately 100 μl of liquid and cell pellet. To lyse the cells, 60 μl lysis buffer (the mixture of solution B and solution C) was added, and the cells were sonicated for 60 cycles (1s on / 2s off) at 30% amplitude using a probe-based sonicator (Q-125, Qsonica, Newtown, CT). The lysed samples were centrifuged, and the resulting supernatant was heated at 99℃ for 1 h. After cooling, solution D was added, mixed thoroughly, and incubated at room temperature for 30 min in the dark. Following this incubation, the samples were centrifuged, and supernatant was transferred into new tubes for protein concentration measurement using the Pierce 660 nm protein assay (IDCR method). Equal amounts of protein were diluted five-fold with solution E, and streptavidin magnetic beads were added into the lysates. The mixture was incubated at room temperature with rotation for 1 h to pull down biotinylated proteins. Protein-bound beads were washed four times using different wash buffers (solution G, H, I and J). The beads were then resuspended in wash buffer and subsequently sonicated for 10 sec in an ultrasonic water bath. The supernatant was carefully discarded, and the beads were resuspended in 20 μl digestion buffer for on-beads digestion. Digestion was carried out at 37℃ for 2 h with shaking using an Intelli-Mixer^TM^ (ELMI, Riga, Latvia). The supernatant was transferred to proteomics tubes provided by the kit and further incubated at 37℃ for 16 h with continuous shaking. The reaction was stopped by solution L, and peptides were desalted. Finally, the desalted peptides were concentrated using a SpeedVac concentrator until completely dried.

Biotinylated proteins were detected using data-independent acquisition (DIA) mass spectrometry. Liquid chromatography-tandem mass spectrometry (LC-MS/MS) was conducted on an UltiMate 3000 RSLCnano system (Thermo Fisher Scientific, Waltham, MA) coupled to an Orbitrap Fusion Lumos mass spectrometer (Thermo Fisher Scientific, Waltham, MA). Desalted and dried peptides were reconstituted in 0.1% formic acid in water and separated on a 75 µm × 250 mm column (pore size 130 Å, particle size 1.7 µm, nanoEase M/Z Peptide CSH C18, Waters). Full MS spectra were acquired using Orbitrap at a resolution of 120,000, with a scan range of 375–1500 m/z, an automatic gain control (AGC) target of 1×10^6^, and a maximum injection time of 50 ms. For DIA MS/MS, 40 scan events were performed with a 10 m/z isolation window, fragmentating ions within the m/z range of 400–800. The MS/MS scan, operating in high-energy collision dissociation (HCD) mode at 30% normalized collision energy, employed an AGC target of 4×10^5^ and a maximum injection time of 54 ms. Fragment ions were scanned in Orbitrap with a resolution of 30,000.

### Mass spectrometry data analysis

Raw DIA files from both labeled and unlabeled (control) samples were analyzed simultaneously using Spectronaut software (version 18.2.230802.50606) in directDIA™ mode, which eliminates the need for a spectral library. Protein identification was conducted using the Pulsar search engine with the UniProtKB/Swiss-Prot human proteome database (Homo sapiens: 20,423 entries) as the reference and default search parameters. Label-free peptide quantification was performed by measuring peak heights at the MS1 level. Protein abundance was estimated by averaging the quantities of the top three most intense peptides, and missing values were imputed using the ‘global imputing’ strategy. After completing protein identification and quantification, the software conducted an unpaired t-test and generated a volcano plot, representing fold changes (labeled/unlabeled) against statistical significance (p-value). Proteins showing at least a 1.5-fold change with a p-value below 0.5 were identified as potential proteins from the regions of interest (ROIs).

### Functional enrichment analysis

Functional enrichment analysis of the identified proteins was performed using Gene ontology (GO) annotation for biological processes (UP_KW_BIOLOGICAL_PROCESS) and cellular components (UP_KW_CELLULAR_COMPONENT) as well as KEGG pathway analysis through the DAVID web server (19). Protein-protein interaction network analysis, including functional clustering, was performed using STRING. Additionally, the Reactome pathway knowledge base was used to map and describe the biological pathways associated with the enriched proteins.

### Image analysis

Confocal image processing and analysis were performed using OpenCV with Python. Signals for mitochondria and lipid droplets were extracted through the following sequential steps:

1. Cell segmentation: All image channels (Tom20, BODIPY 558/568 and DAPI) were merged to define cell boundaries. The merged images were converted into binary format using the watershed algorithm.
2. Target extraction: Mitochondria or lipid droplet images were extracted using an adaptive thresholding algorithm. A size filter was applied to exclude small noise particles, producing clear binary images of both targets.
3. The ‘AND’ logical operation was used to confirm that both target binary images were within the segmented cell image.

To quantify mitochondria-LD contacts, the ‘AND’ function was applied to identify co-localization pixels of mitochondria and lipid droplets, and the contact numbers were calculated. For measuring mitochondria aspect ratio measurement, isolated mitochondria were identified using contour extraction. The minimum bounding rectangle of each contour was determined, allowing the calculation of the width-to-height ratio. LD size was assessed by analyzing extracted LD image signals using the ‘Analyze Particles’ function in ImageJ software to determine their area.

### Western blot

HepG2 cells were washed once with ice-cold PBS and then added modified RIPA buffer (50 mM Tris-HCl pH7, 150 mM NaCl, 1 mM EDTA, 1% Triton X-100, 0.5% SDS and 1x protease inhibitor cocktail) to scrap and harvest cells for 20 min on ice. DNA was sheared, and cell disruption was performed using sonication (30% power, 6 pulses). The lysates were centrifuged at 4℃ for 10 min, and the supernatant was collected. Protein concentration was measured using Bio-Rad Bradford method, and equal amounts of protein were denatured at 95℃ for 5 min before separation by SDS-PAGE. Proteins were transferred onto PVDF membrane, followed by blocking with blocking solution (BF01, Bio Pioneer Tech, Taiwan) and washing with PBS. The membranes were incubated overnight at 4℃ with either mouse anti-α-tubulin or rabbit anti-FHL3 antibodies. Subsequently, they were incubated for 1 hour at room temperature with HRP-conjugated goat anti-mouse or rabbit antibodies. After three washes with PBS, signals were detected using ECL subtract (#1705061, Bio-Rad Laboratories, CA) and visualized using the iBright FL1500 image system (Thermo Fisher Scientific, Waltham, MA).

### Statistical analysis

Immunoblot band intensities was quantified using imageJ software, and statistical analyses were performed using GraphPad Prism 10 software (GraphPad, San Diego, CA). Data were presented as the mean ± SD from at least three independent experiments. Statistical significance between two groups was assessed using Welch’s t-test, while one-way ANOVA followed by Tukey’s post hoc test was used for multiple group comparisons. A p-values of less than 0.05 was considered statistically significant.

## Results

### Defining and photolabeling mitochondria-LDs contact sites using Microscoop technology

Microscoop Mint is an advanced platform that integrates image acquisition, photochemistry, microscopy and optical system to enable high content in situ biotinylation of proteins at specified cellular regions with subcellular resolution. When coupled with pulldown and mass spectrometry analysis, Microscoop facilitates the precise identification of proteins localized to region of interest. The Microscoop to LC-MS/MS workflow consists of the following steps (Fig. 1A):

1. Preparation of cell samples and visualization of the ROIs: Cells are prepared, and the regions of interest (ROIs) are detected using corresponding antibodies or through the ectopic expression of fusion protein tagged with fluorescence proteins. Photo-activatable biotin probes are added to the samples.
2. Imaging and photolabeling: Image is first acquired by the Microscoop Mint on a single of view (FOV). ROIs are then recognized using image processing functions which define the ROI and generate mask patterns to guide two-photon laser movement for photolabeling. The system automatically cycles through all selected FOVs. Photolabeling is completed until all FOVs are processed.
3. Protein extraction and identification: Biotinylated proteins are extracted, purified, and identified via LC-MS/MS.

**Figure 1.**
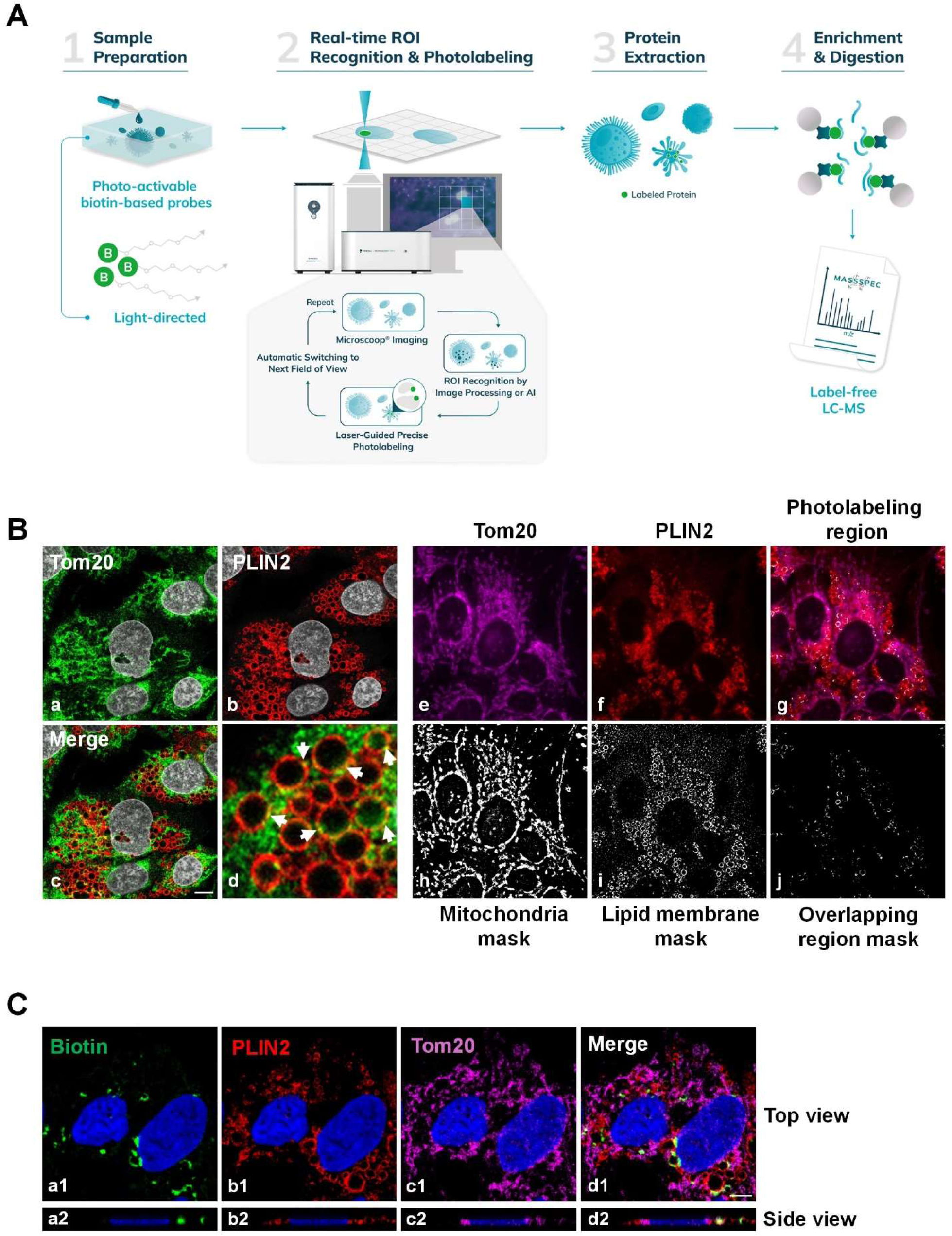
ROI recognition and photolabeling of mitochondria-LD contact sites using Microscoop Mint platform. (A) Schematic workflow of Syncell Microscoop Mint. A total-sync ultra-content microscopic platform that integrates image acquisition, photochemistry, microscopy, optics, and FPGA-based mechatronics to enable high-content in situ photolabeling followed by mass spectrometry analysis. (B) Left panel: Immunostaining of mitochondria and LD membranes using Tom20 (a, green) and PLIN2 (b, red), respectively. The overlapping regions (c and d) of Tom20 and PLIN2 signals indicate potential mitochondria-LD contact sites. Right panel: Mitochondria (e) and LD (f) images (e and f) are recognized using traditional image processing and computer vision algorithm (h and i) to identify ROIs (g and j, mitochondria-LD overlap region). (C) Confocal images demonstrate the precise and accurate photo-biotinylated proteins (a1) within mitochondria-LD contact regions in both lateral (xy)- and axial (z) directions (a1-d1 and a2-d2). Scale bar: 5 μm.

To visualize mitochondria-LD contact sites, lipid droplet accumulation in HepG2 cells was induced with oleic acid (OA) treatment. Mitochondria and LDs were immuno-stained with antibodies against Tom20 (mitochondria outer membrane; Fig. 1B, a) and PLIN2 (LD membrane; Fig. 1B, b), respectively. The merged image (Fig. 1B, c and d) revealed overlapping regions of Tom20 and PLIN2 signals, marked by yellow, indicating the potential mitochondria-LD contact sites. Using Microscoop Mint, images were first acquired to generate mask patterns for mitochondria (Fig. 1B, e and h) and LD membrane (Fig. 1B, f and i) through traditional image processing functions (described in Materials and Methods). The overlapping regions were then defined as masks (Fig. 1B, g and j) for targeted photolabeling. Following photolabeling, the localization of biotinylated proteins was verified using confocal imaging, which confirmed their precise localization within the mitochondria-LD overlapping regions in both lateral (xy) and axial (z) directions (Fig. 1C). This indicates that Microscoop enables highly accurate and specific in situ spatial photolabeling.

### Spatial proteomic characterization of mitochondria-LD contact region

To characterize the proteome of the mitochondria-LD contact site, photolabeled and unlabeled HepG2 cells were harvested, and biotinylated proteins were isolated and identified via mass spectrometry. The LC-MS/MS dataset, conducted in three independent replicates, was filtered using the criteria of log_2_ (fold change) ≧ 0.58, -log (p-value) ≧ 1.3, and unique peptide ≧ 2. This analysis identified a total of 2,264 enriched proteins in the photolabeled group compared to the unlabeled control (Supplementary Table 2). The dataset was cross-referenced with the mitochondria protein databases (Uniprot and human MitoCarta 3.0) and a curated database of 231 known LD-localized proteins (compiled from Uniprot and prior literature supported by immunofluorescence and mass spectrometry evidence) (20, 21). The comparison revealed that 75 high-confident well-known LD-localized proteins and 561 mitochondrial proteins were enriched in the photolabeled group (Fig. 2A). These LD-localized and mitochondrial proteins (636 in total) were subjected to Gene Ontology (GO) analysis, which revealed significant enrichment in biological processes such as the tricarboxylic acid cycle (TCA), fatty acid metabolism, and lipid biosynthesis and metabolism (Fig. 2B, left panel, Supplementary Table 3).

**Figure 2.**
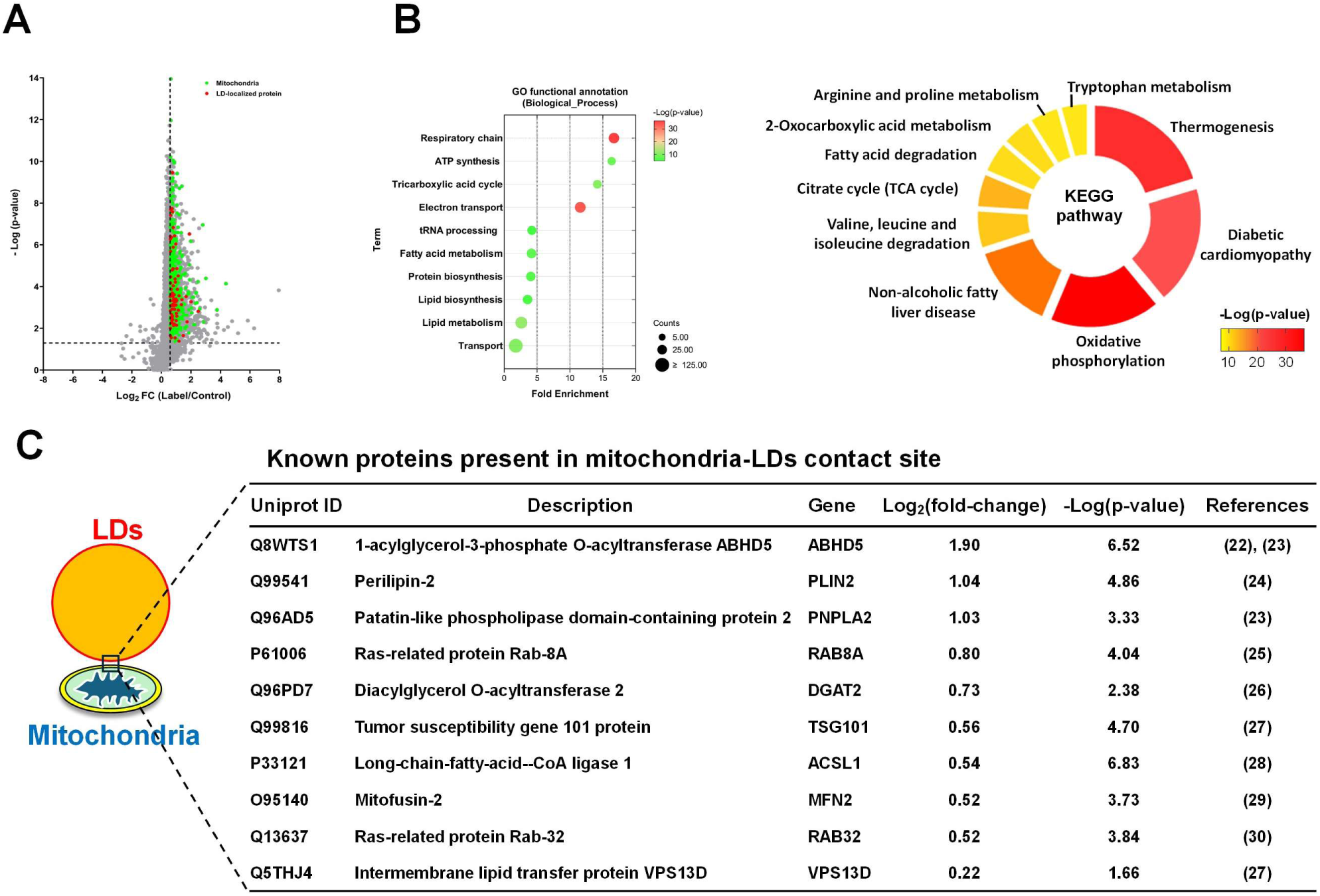
Spatial proteomics analysis of mitochondria-LD contact sites. (A) Distribution of overall protein abundances represented as the ratio of photolabeled (PL) samples to control (CTL) samples, expressed as PL/CTL ratio. Known LD-localized protein (blue) and mitochondria proteins (red) are significantly enriched in the PL group. (B) Gene Ontology (GO) enrichment analysis of biological processes was performed on a combined dataset of 75 known LD-associated proteins and 561 known mitochondria proteins (left panel). KEGG pathway analysis (right panel) illustrates the distinct pathways associated with these proteins. (C) The list of the known proteins involved in mitochondria-LD interactions identified using the Microscoop Mint platform.

KEGG pathway analysis further highlighted significant enrichments in pathways such as TCA cycle, fatty acid degradation, and NAFLD (Fig. 2B, right panel), underscoring the relevance of these proteins to lipid-related functional processes and NAFLD pathogenesis. Notably, proteins previously reported to mediate mitochondria-LD association were identified in the dataset (Fig. 2C) (22–30). They include tumor susceptibility gene 101 protein (TSG101), a non-LD and non-mitochondria protein recruited by VPS13D to mediate the fatty acid transfer at mitochondria-LD contact site (27). These findings validate the reliability, sensitivity and specificity of Microscoop photolabeling in identifying proteins at mitochondria-LD contact regions and provide new insights into lipid-related functional processes, including NAFLD.

To identify novel proteins potentially located at the mitochondria-LD contact site, we analyzed the filtered datasets from three independent proteomic experiments using stringent criteria: log_2_ (fold change) ≧ 0.58, -log (p-value) ≧ 1.3, and unique peptide ≧ 2. Among the enriched proteins, 373 were consistently identified across all three replicates (Fig. 3A, Supplementary Table 4). Gene Ontology (GO) analysis was performed on these 373 proteins to assign cellular component terms. While a large portion of the proteins were categorized as associated with mitochondria or LDs, a subset was linked to the endoplasmic reticulum (ER) and peroxisomes (Fig. 3B, left panel). GO biological process analysis of the top 10 enriched terms revealed that only 7.5% of these proteins were directly annotated for lipid metabolism, while the majority were involved in diverse functional categories (Fig. 3B, right panel). This suggests that many of these high-confidence proteins may have indirect or previously uncharacterized roles in lipid-related processes.

**Figure 3.**
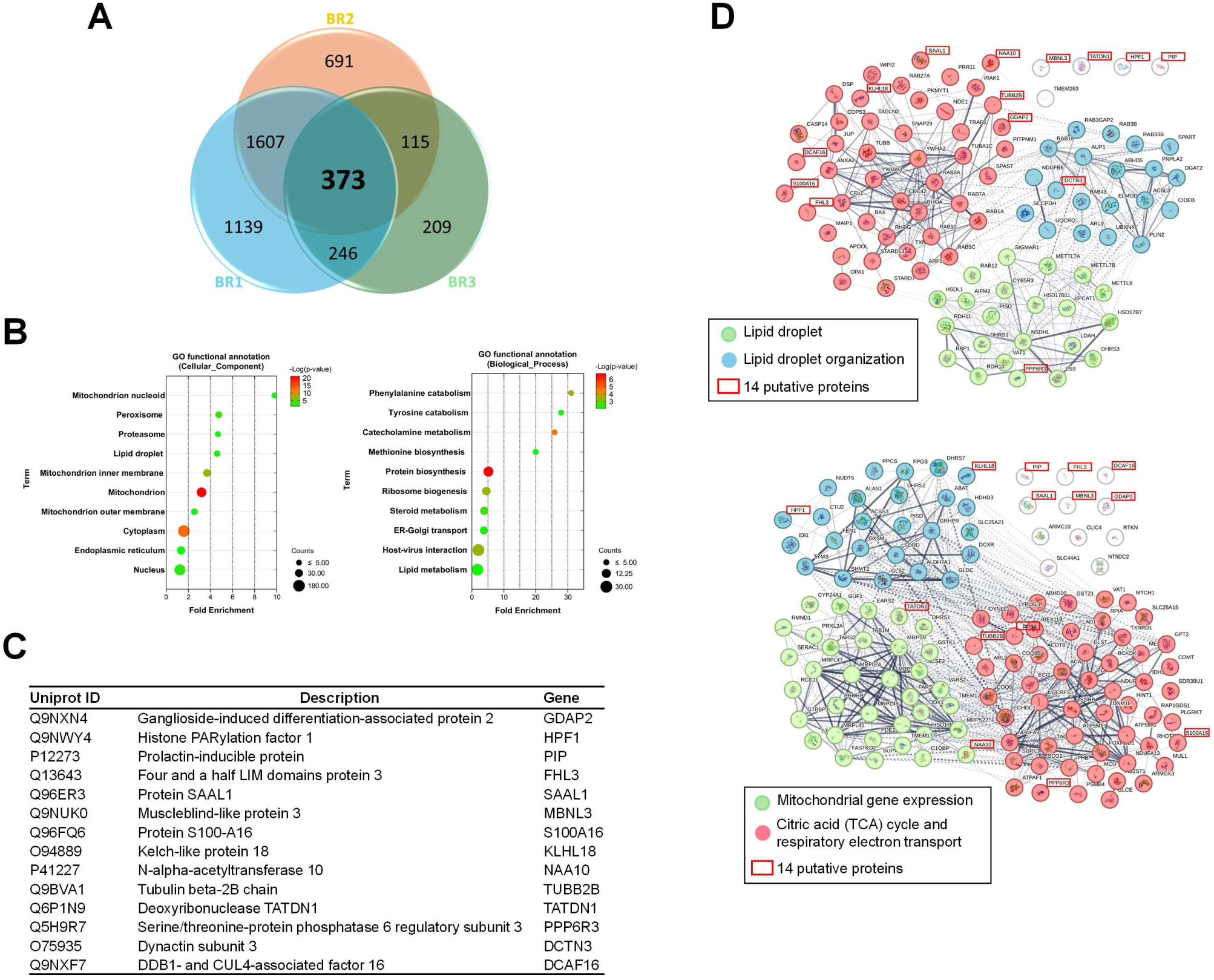
Functional enrichment analysis of identified putative proteins. (A) Venn diagram depicting 373 common proteins consistently identified across three replicate experiments. (B) Functional enrichment analysis of these 373 common proteins, highlighting the top 10 enriched GO Cellular Component terms and the top 10 functional annotation clusters of Biological Process terms. (C) The list of 14 putative proteins selected from the top 30 enriched common proteins. (D) STRING analysis of 14 putative proteins combined with enriched known LD-associated proteins (upper panel) or mitochondria proteins (lower panel) identified using the Microscoop Mint platform to assess protein-protein interaction relationships. Proteins with similar functional processes are represented in clusters of the same color.

To uncover potential candidates, we selected 14 proteins (Fig. 3C) from the top 30 common proteins (Table 1) that were not previously annotated as mitochondria or LD associated proteins. STRING analysis was performed to evaluate protein-protein interaction (PPI) networks involving these 14 proteins alongside enriched mitochondria proteins (111 proteins) and LD-associated proteins (75 proteins) to deduce potential interactions. Interestingly, PPI analysis showed that these 14 proteins are more closely related to the functional cluster of LD-associated proteins (Fig. 3D upper panel) than to mitochondria proteins (Fig. 3D lower panel), suggesting their potential involvement in LD-related processes. Further GO “Cellular Component” analysis of the 14 candidates revealed subcellular localization such as the cytoplasm, nucleus, lysosome, extracellular exosome, and histones (Supplementary Table 5). To narrow down candidates for immunostaining validation, we prioritized proteins with the highest fold change and excluded those associated with lysosomes, histones or exosomes. This refinement resulted in five candidates: four proteins with no prior association to lipid metabolism or mitochondria, including four-and-a-half LIM domains protein 3 (FHL3), protein SAAL1 (SAAL1), muscleblind-like protein 3 (MBNL3) and N-alpha-acetyltransferase 10 (NAA10), along with armadillo repeat-containing protein 10 (ARMC10), a mitochondria protein with the highest fold change enrichment among the 373 common proteins.

**Table 1.**
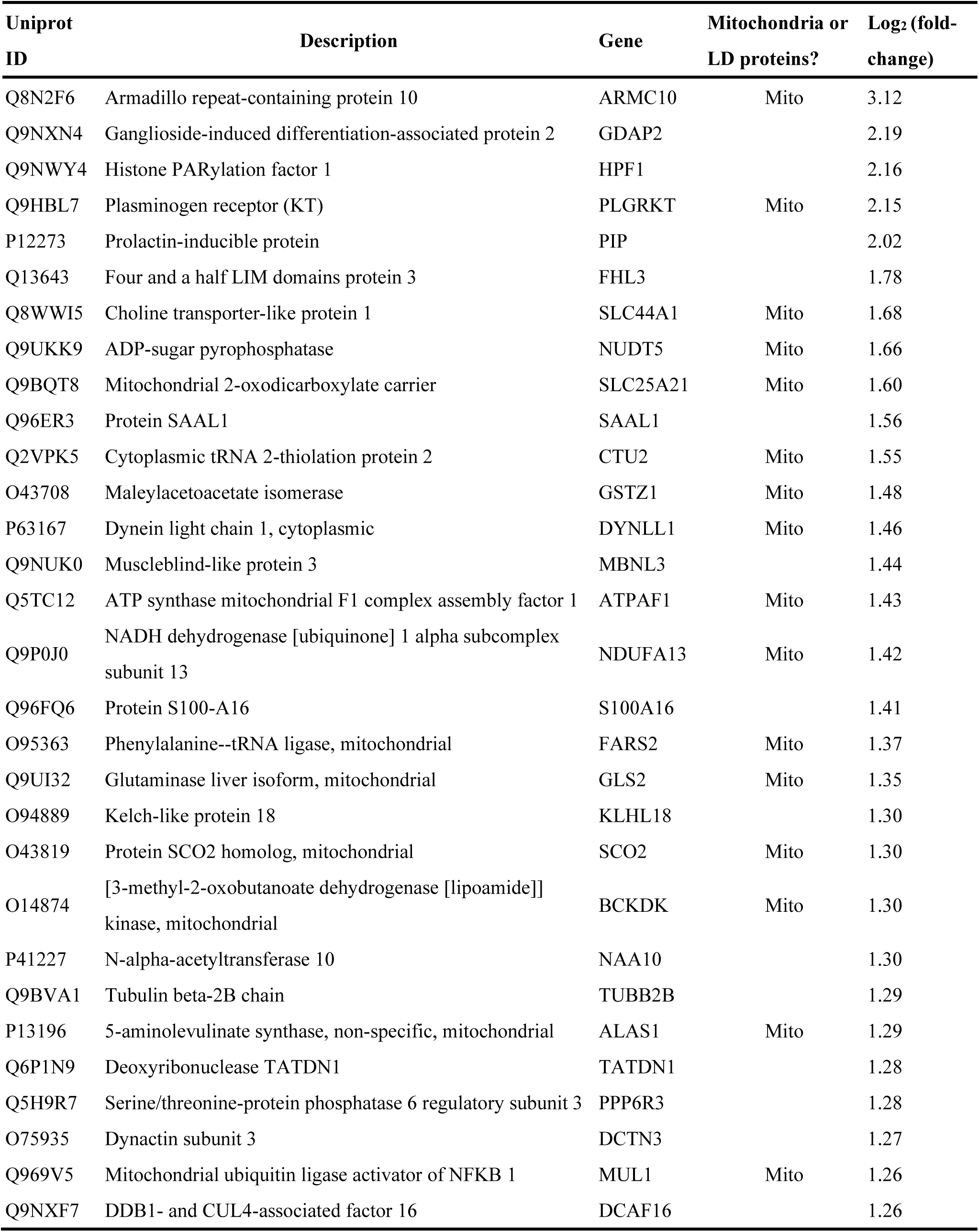
List of Top 30 common proteins.

### The putative proteins were associated with LDs and localized at mitochondria-LD contact sites

Previous studies have suggested that certain proteins bind to LDs via amphipathic helices domain within their protein sequences, contributing to LDs homeostasis or mediating LD-mitochondria communication (27, 31, 32). Amphipathic helices are α-helix structures in proteins characterized by hydrophobic and hydrophilic faces arranged on opposite sides, formed by hydrophobic and polar amino acid residues. This topology facilitates the efficient adsorption of such proteins onto the LD surface. To determine whether the five candidates contain potential amphipathic helices, we first conducted secondary structure predictions of their sequences using three algorithms (PHD, SOPM and SOPMA) on the NPS@ (Network Protein Sequence Analysis) website (33). Predicted α-helix regions were further analyzed using HeliQuest program (34) to assess their amphipathic properties. The results demonstrated that all five proteins possess potential amphipathic α-helix regions, suggesting that the hydrophobic face of these helices may facilitate their association with LD surfaces (Fig. 4A). Based on these predictions, we hypothesized that these five proteins might associate with LDs or lipid surfaces of cellular organelles. To test this hypothesis, we examined whether these five candidate proteins co-localize with LDs and are present in mitochondria-LD contact sites. Epi-fluorescence imaging showed that all five proteins localized around LDs, with more pronounced signals in OA-treated HepG2 cells compared to vehicle control (Fig. 4B). Representative confocal images further confirmed their colocalization at LD membranes, displaying distinct patterns: a dot-like distribution (ARMC10 and MBNL3; Fig. 4C, a1-a3 and d1-d3) and a crescent-like arrangement (FHL3, SAAL1 and NAA10; Fig. 4C, b1-b3, c1-c3 and e1-e3) at the LD periphery, with some forming a ring-like structure (FHL3, Fig. 4, b1-b3) encircling the LDs. Additionally, these proteins were observed at mitochondria-LD contact sites (Fig. 4C, merged images of a6, b6, c6, d6 and e6). This localization suggests a potential role for these novel proteins in regulating mitochondria-LD interaction or maintaining LD homeostasis.

**Figure 4.**
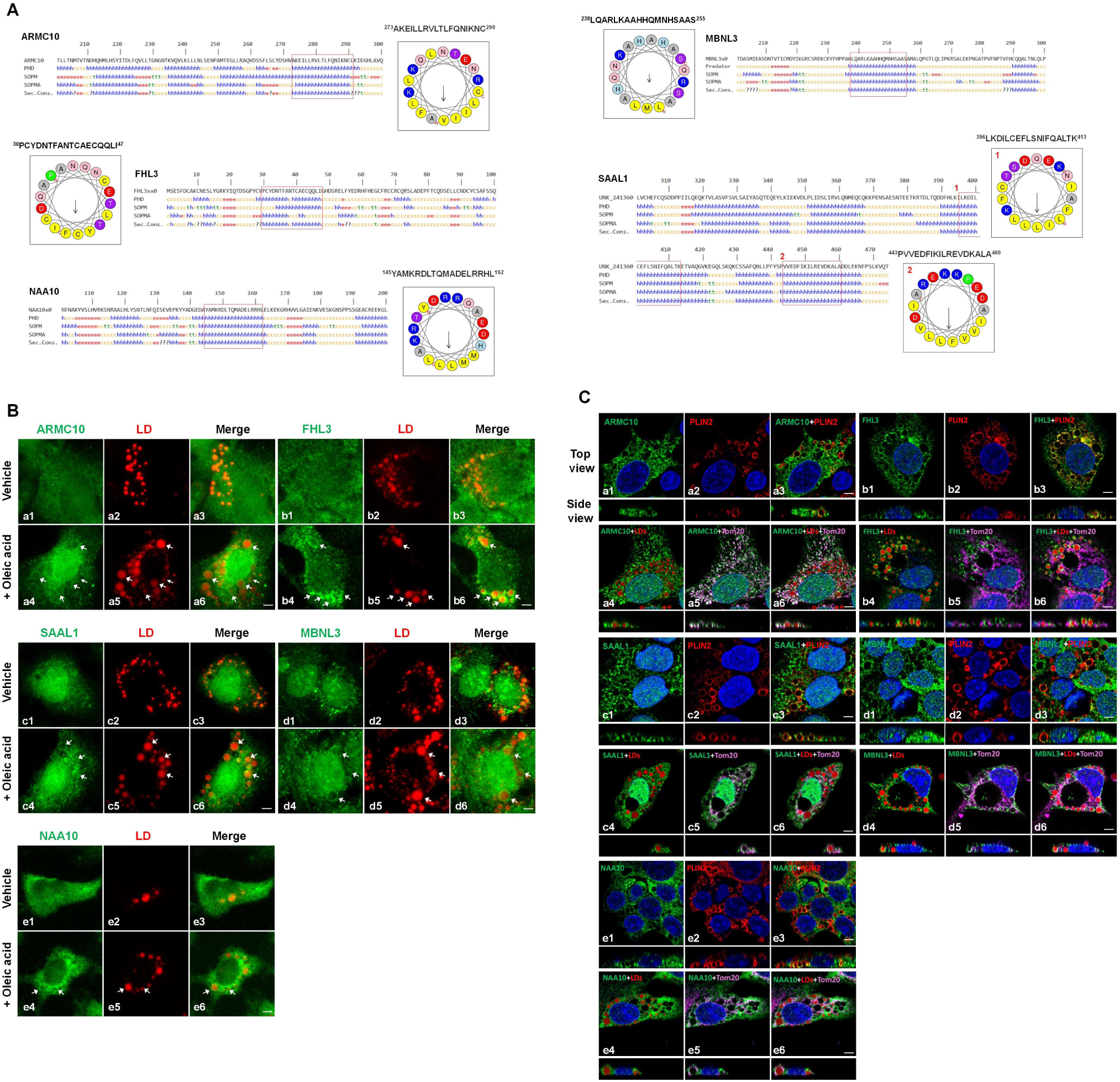
Discovery of novel proteins associated with mitochondria-LDs contact sites. (A) Secondary structure and amphipathic helix prediction of 5 putative LD-associated proteins, selected from the Top 30 common proteins, using algorithms from the NPS@ website and HeliQuest program respectively. The analysis highlights potential amphipathic α-helix regions within these 5 proteins. (B) Lipid droplets in oleic acid-treated HepG2 cells were stained with BODIPY 558/568 (red), and the 5 candidate proteins were detected using corresponding antibodies. Epi-fluorescence images revealed their localization around lipid droplets (arrows), with more prominent association in OA-treated HepG2 cells compared to vehicle-treated controls. Scale bar: 5 μm. (C) Confocal images verified the co-localization of all 5 candidate proteins with lipid droplet membrane (PLIN2 as a marker), as well as their presence at mitochondria-LD contact sites, suggesting a potential implication of these novel proteins in mitochondria-LD regulation. Scale bar: 5 μm.

### FHL3 deficiency alters mitochondria morphology and reduces mitochondria-LD interaction

To gain biological insights in the role of the five candidates, we conducted a Reactome pathway analysis integrated with ‘IntAct’ molecular interaction database. Notably, FHL3 was suggested to be implicated in mitochondrial fatty acid β-oxidation within the fatty acid metabolism pathway, potentially along with its interaction partner, NADH-ubiquinone oxidoreductase subunit AB1 (NDUFAB1) (Fig. 5A). NDUFAB1, a mitochondrial protein with acyl carrier activity and fatty acid binding functions, was previously identified as a FHL3 interactor via a yeast two-hybrid assay (35). Previous reports have shown that NDUFAB1 participates in fatty acid biosynthesis through interactions with other proteins (36) and may also play a role in fatty acid β-oxidation in fungi (37). Based on these findings, we selected FHL3 for further functional studies.

**Figure 5.**
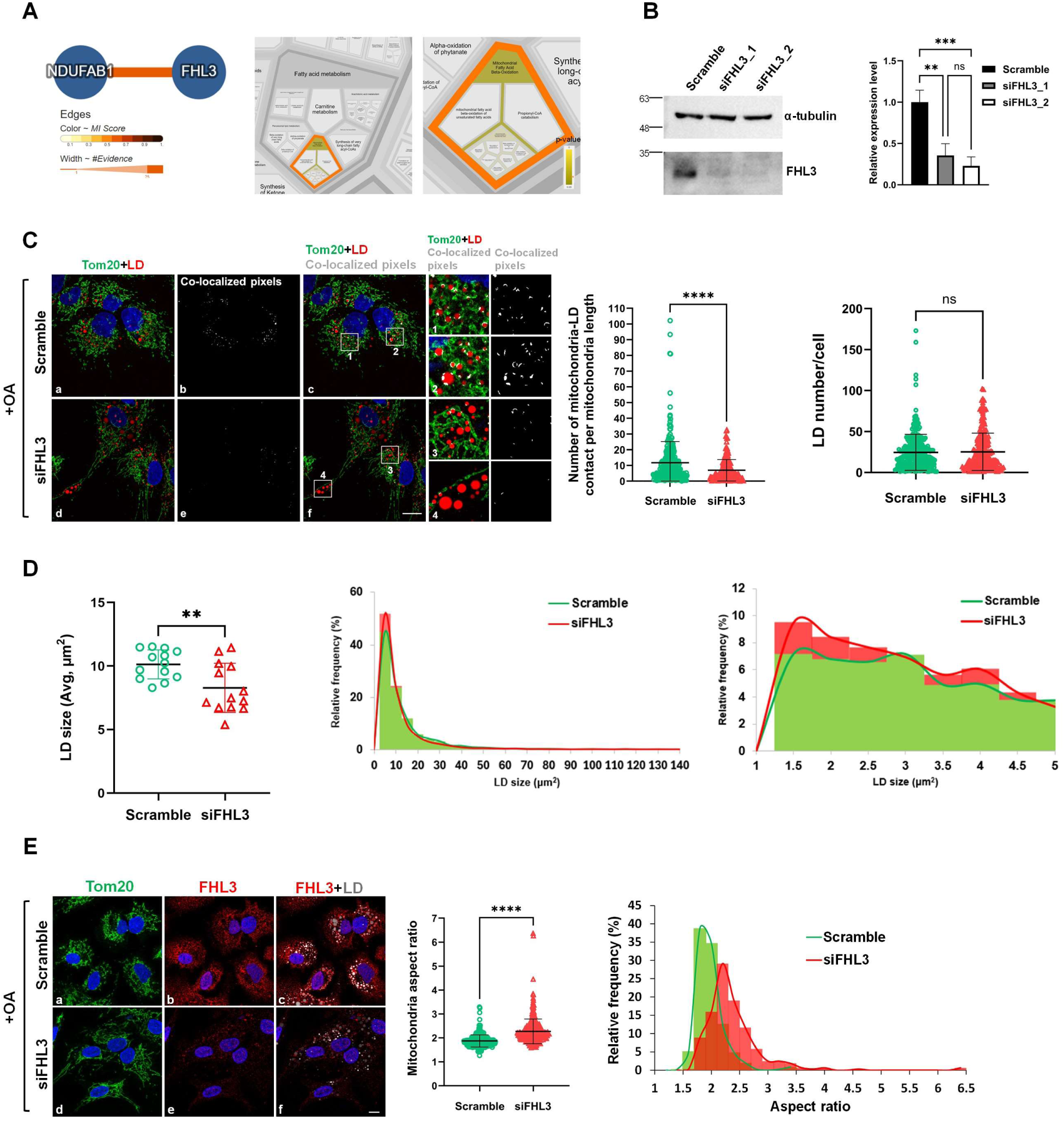
Mitochondria morphology alteration and reduction of mitochondria-LD interaction upon FHL3 depletion. (A) Reactome analysis suggests that FHL3 may be involved in mitochondrial fatty acid β-oxidation through its potential interaction with NDUFAB1. (B) Western blot demonstrated the expression level of FHL3 after siRNA transfection (left panel), with quantification confirming a significant decrease in FHL3 protein in siRNA-treated HepG2 cells (right panel, **, p < 0.01, ***, p < 0.001, ns: not significant). α-tubulin was used as a loading control. (C) Confocal images of scramble control (a-c) or siFHL3-transfected (d-f) HepG2 cells stained for Tom20 (green) and BODIPY 558/568 (red) upon OA treatment. The white spots represent the co-localized pixels of mitochondria-LD contact site (b and e). Enlarged views of the boxed regions in the merged images (c and f) are shown on the right. The quantification result of the number of mitochondria-LD contact per mitochondria length (middle panel) and the lipid droplet number per cell (right panel) were calculated in scramble (total 270 or 289 cells respectively) or siFHL3 (total 204 or 231 cells respectively)-treated HepG2 cells from four independent experiments (****, p < 0.0001, ns: not significant). (D) The LDs area of OA-treated scramble or FHL3-depleted HepG2 cells were measured from three independent experiments (left panel; measured from 229 and 177 cells for control and FHL3-depleted cells respectively; **, p < 0.01). The overall distribution of LDs size in scramble or siFHL3-treated cells was shown in the middle panel, while the distribution of LDs size smaller than 5 μm^2^ was presented in the right panel. (E) The representative confocal images of elongated mitochondria morphology in FHL3 deficiency HepG2 cells (d-f) compared to scramble control (a-c). The quantification scatter plot of mitochondrial aspect ratio of scramble (total 348 cells) and siFHL3 cells (total 326 cells) were calculated from four independent experiments (middle panel), with the aspect ratio distribution shown in the right panel (****, p < 0.0001). Scale bar, 10 μm in whole cell images.

To investigate FHL3’s role in mitochondria-LD interactions, we knocked down its expression using RNA interference. Western blot analysis confirmed efficient knockdown of FHL3 protein levels in HepG2 cells transfected with two different FHL3 siRNAs (siFHL3_1 and siFHL3_2) compared to a scramble siRNA control (Fig. 5B). We examined the effect of FHL3 depletion on mitochondria-LD interaction by measuring the co-localization of Tom20 and PLIN2 with fluorescence microscopy (Fig. 5C, left panel). FHL3 knockdown resulted in a ∼40% reduction in the number of co-localized pixels between mitochondria and LDs (Fig. 5C, middle panel), indicating impaired mitochondria-LD interactions, though the average number of LDs were not significantly affected (Fig. 5C, right panel). Interestingly, FHL3 deficient cells showed a significant reduction in LD size (Fig. 5D, left panel). Summarizing distribution of LD size revealed an increased proportion of small LDs (< 5 μm²) in FHL3-depleted cells, with a 6.8% increase as compared to the scramble control (Fig. 5D, middle and right panels). These results suggest that disrupting mitochondria-LD interactions impairs LD growth under OA overload conditions.

Tom20 immunofluorescence staining further revealed that the mitochondria in FHL3-deficient, OA-treated HepG2 cells exhibited elongated morphology compared to the fragmented mitochondria observed in scramble control cells (Fig. 5E, left panel). Quantitative analysis showed a significantly higher mitochondrial aspect ratio (long axis to short axis) in FHL3-deficient cells compared to controls (Fig. 5E, middle panel). Distribution analysis indicated that 74.8% of mitochondria in FHL3-deficient cells had an aspect ratio > 2, compared to only 20.9% in control cells (Fig. 5E, right panel). Since elongated mitochondria have been shown to exhibit reduced fatty acid β-oxidation rates compared to the fragmented mitochondria (18), we speculated that FHL3 suppression disrupts mitochondria-LD interactions, leading to elongated mitochondria and attenuated fatty acid β-oxidation, ultimately affecting lipid metabolism.

In summary, using Microscoop coupled with LC-MS/MS, we successfully isolated and identified proteins enriched at the mitochondria-LD contact sites. They include known mitochondria and LD-associated proteins, previously reported mediators of mitochondria-LD interactions, and novel proteins with no prior LD association. Five putative proteins were validated by immunostaining and found to be localized at mitochondria-LD contact sites. Further biological studies of one of the proteins, FHL3 revealed its potential role in lipid metabolism. FHL3 deficiency significantly diminished mitochondria-LD interactions. In addition, depletion of FHL3 reduced LD size and induced mitochondrial elongation in OA-treated HepG2 cells. These findings suggest a critical role of FHL3 in mediating mitochondria-LD interactions and regulating lipid metabolism.

## Discussion

Intracellular organelles are membrane-bound structures that compartmentalize various biological processes, enabling incompatible reactions to occur simultaneously without interference. Beyond studying the functional mechanisms, protein composition, and unique characteristics of individual organelles, our research focuses on the interactions between intracellular organelles. Proteins within membrane contact sites (MCSs) are widely recognized as key mediators of physical contact and functional coordination between organelles. These interactions play critical roles in regulating organelle morphology, signal, and metabolite transfer, allowing cells to respond effectively to various stimuli (1). A prominent example is the bi-directional transfer of fatty acids at mitochondria-LD contact sites, which supports lipid metabolism or storage. This process is essential for energy production during nutrient scarcity (4), and for mitigating lipotoxicity or oxidation stress caused by excess fatty acid accumulation (38). Disruptions in mitochondria-LD interactions have been linked to a range of diseases, including neurological disorders, cancer and metabolic diseases such as type 2 diabetes and NAFLD (39). Identifying and investigating the biological functions of novel proteins involved in mitochondria-LD interactions or other organelles’ cross talk is therefore crucial. Such discoveries could provide deeper our understanding of these cellular processes and insights into these potential therapeutic targets for treating related diseases.

Given the critical biological function of mitochondria-LD interactions, studying the inter-organelle contact proteome is both intriguing and challenging due to the complicated and dynamic nature of these interactions and the lack of efficient tools. While biochemical methods for isolating LDs followed by mass spectrometry analysis are well established, prior studies have shown that mitochondria often co-isolated with LDs and can be observed on the surfaces of isolated spherical LDs (40). However, in certain cell types, some LD-anchored mitochondria may detach under higher centrifugal forces, while others remain tightly associated (41). This highlights the risk of proteins being stripped from mitochondria-LD membrane contact sites during the purification process, particularly those involved in transient, dynamic connections via ‘kiss-and-run’ model. Moreover, purification protocols often need to be adjusted and optimized for different cell or tissue types, introducing variability and uncertainty in experimental results. To address these limitations, we employed the Microscoop Mint system, a microscopy-guide spatial protein isolation platform with submicron precision, to investigate the protein composition of mitochondria-LD contact sites and identify novel proteins involved in these interactions under OA-treatment conditions.

Using mitochondria and LD membrane markers for visualization, real-time image analysis and processing enabled segmentation of mitochondria-LD contact regions. The platform efficiently segmented each FOV image of the contact site and used two-photon illumination to trigger photoactivation of biotinylation reagents, specifically labeling proteins localized to the contact sites. Mass spectrometry analysis of the labeled regions identified numerous known LD-localized and mitochondria proteins, along with several previously reported regulators of mitochondria-LD interactions (Fig. 2C). From our proteomic analysis, we selected five putative proteins from the top 30 ranked by fold change for further validation. Immunofluorescence staining demonstrated that all five proteins colocalized to the LD membrane and were present at mitochondria-LD contact sites in OA-treated HepG2 cells, suggesting their potential involvement in regulating mitochondria-LD interactions. This innovative approach overcomes the limitations of traditional methods and serves as a robust platform to uncover proteins critical to inter-organelle communication.

In recent years, proximity labeling technology has emerged as a powerful tool for identifying proteins in close spatial proximity to a target protein within living cells. This approach typically involves the ectopic expression of genetically fused specific protein or subcellular compartment markers with labeling enzymes, such as peroxidase (APEX and APEX2) or biotin ligase (BioID2, TurboID2 and BASU etc.). The labeling enzymes react with biotin substrates, resulting in the biotinylation of nearby proteins, which can then be identified through mass spectrometry to study protein-protein interactions or the proteomes of specific subcellular compartments (42). Compared to Microscoop, proximity labeling offers advantages such as higher spatial resolution (∼20 nm labeling radius), shorter labeling times (particularly with peroxidase enzymes), and the ability to label both proteins and RNA. However, this approach requires the construction of genetic fusion constructs for the labeling enzymes and their ectopic expression in cells, often achieved through transfection or virus induction. These overexpressed fusion proteins can potentially alter the location, function, or interactome of the target proteins or subcellular compartments. Additionally, peroxidase-base labeling relies on hydrogen peroxide to catalyze the reaction, which can disrupt cellular redox pathways and poses toxicity concerns, limiting its use in vivo.

The dependence on exogenous fusion construct expression presents further challenges for applying proximity labeling to primary or clinical human tissue samples. To address this limitation, a previous study developed an antibody-guided proximity labeling method, which used an HRP-conjugated secondary antibody in combination with phenol biotin and hydrogen peroxide to biotinylate proteins near antibody-recognized targets (43). While this approach avoids genetic manipulation, it is constrained by non-specific protein biotinylation, which can interfere with downstream proteomics analysis. Furthermore, its reliance on highly specific antibodies that perform well in tissue sections limits its applicability in human tissue samples. In contrast, Microscoop offers a distinct advantage through its image-guided target recognition and use of a photoactivatable biotin labeling probe. This probe is activated by two-photon focal light, enabling highly specific and precise labeling of proteins within defined image mask regions. This approach is adaptable to a variety of sample types, including cells, mouse tissues, and human clinical tissue sections, providing broader applicability while maintaining specificity and reducing background noise.

Although the precise etiological link between mitochondria-LD interactions and NAFLD remains incompletely understood, it is well established that proteins involved in these interactions play a crucial role in lipid metabolism and mitochondrial integrity (13). Dysregulation of either process can contribute to the development and progression of NAFLD. Therefore, identifying novel proteins associated with mitochondria-LD interactions and conducting proteome analyses of mitochondria-LD contact sites under different disease condition models could provide valuable insights for understanding NAFLD pathophysiology and facilitate therapeutic advancements. Using Microscoop Mint platform, we identified and validated five novel LD-associated proteins potentially involved in the regulation of mitochondria-LD functional interactions. RNA interference experiments targeting one of these proteins, FHL3, demonstrated that its depletion led to mitochondria elongation, a significant reduction in mitochondria-LD contacts, and smaller LD size in OA-treated HepG2 cells. These findings suggest that FHL3 depletion may disrupt the balance between mitochondria-LD interactions and LD growth under OA overload. Moreover, the observed elongated mitochondrial elongation is indicative of reduced β-oxidation activity (18), a phenomenon that has also been reported in NAFLD rat liver models (40). Based on these findings, we hypothesize that FHL3, identified from mitochondria-LD contact sites through the Microscoop to LC-MS/MS workflow, may play a critical role in maintaining mitochondria-LD interactions and regulating lipid metabolism, potentially contributing to NAFLD pathogenesis. The Microscoop Mint platform not only enables advances in proteomic research but also facilitates the identification of novel potential drug targets across diverse disease models for therapeutic advancements.

## Acknowledgements

We thank the Core Facilities of Translational Medicine of BioTReC (National Biotechnology Research Park, Academic Sinica), Taiwan for the technical support and data analysis, and Professor Hsueh-Fen Juan at National Taiwan University for the useful discussion.

## Declaration of competing interest

Patent applications related to the subject matter of this publication have been filed. All authors declare that they are current employees of Syncell Inc., and, except for YML, they are also shareholders in the company.

