## Supplementary material for "Microscopy-Guided Spatial Proteomics Reveals Novel Proteins at the Mitochondria-Lipid Droplet Interface and Their Role in Lipid Metabolism": Supplememtary Table 1.docx

Supplementary Table 1

| Antibodies | Source | Identifier |
| --- | --- | --- |
| Rabbit polyclonal anti-PLIN2 | Proteintech | Cat# 15294-1-AP,  RRID: AB_2878122 |
| Mouse monoclonal anti-PLIN2 | Santa Cruz Biotechnology | Cat# sc-377429  RRID: AB_3661746 |
| Mouse monoclonal anti-Tom20 | Santa Cruz Biotechnology | Cat# sc-17764,  RRID: AB_628381 |
| Rabbit recombinant monoclonal anti-Tomm20 | Abcam | Cat# ab186735,  RRID: AB_2889972 |
| Rabbit polyclonal anti-ARMC10 | Proteintech | Cat# 20506-1-AP,  RRID: AB_10696176 |
| Rabbit polyclonal anti-SAAL1 | Proteintech | Cat# 25467-1-AP,  RRID: AB_2880094 |
| Rabbit polyclonal anti-MBNL3 | Proteintech | Cat# 24610-1-AP,  RRID: AB_2879637 |
| Mouse monoclonal anti-NAA10 | Santa Cruz Biotechnology | Cat# sc-373920,  RRID: AB_10918374 |
| Mouse monoclonal anti-FHL3 | Santa Cruz Biotechnology | Cat# sc-166917,  RRID: AB_10612720 |
| Rabbit polyclonal anti-FHL3 | Proteintech | Cat# 11028-2-AP,  RRID: AB_2294066 |
| Mouse monoclonal anti-α-tubulin | GeneTex | Cat# GTX11304,  RRID: AB_373242 |
| Donkey anti-Rabbit IgG,  Alexa Fluor 488 | Thermo Fisher Scientific | Cat# A-21206,  RRID: AB_2535792 |
| Donkey Anti-Mouse IgG,  Alexa Fluor 488 | Thermo Fisher Scientific | Cat# A-21202,  RRID: AB_141607 |
| Goat anti-Rabbit IgG,  Alexa Fluor 568 | Thermo Fisher Scientific | Cat# A-11011,  RRID: AB_14315 |
| Donkey Anti-Mouse IgG,  Alexa Fluor 568 | Thermo Fisher Scientific | Cat# A10037,  RRID: AB_11180865 |
| Goat anti-mouse IgG,  Alexa Fluor 647 | Thermo Fisher Scientific | Cat# A-21235,  RRID: AB_2535804 |
| Goat anti-Rabbit IgG,  Alexa Fluor 647 | Thermo Fisher Scientific | Cat# A-21245,  RRID: AB_2535813 |
| Goat Anti-Mouse IgG, HRP | GeneTex | GTX213111-01,  RRID: AB_10618076 |
| Goat Anti-rabbit IgG, Peroxidase-Labeled | SeraCare KPL | Cat# 5450-0010,  RRID: AB_3075498 |
